# The impact of rheotaxis and flow on the aggregation of organisms

**DOI:** 10.1101/2021.07.20.453166

**Authors:** Kevin J. Painter

## Abstract

Dispersed populations often need to organise into groups. Chemical attractants provide one means of directing individuals into an aggregate, but whether these structures emerge can depend on various factors, such as there being a sufficiently large population or the response to the attractant being sufficiently sensitive. In an aquatic environment, fluid flow may heavily impact on population distribution and many aquatic organisms adopt a rheotaxis response when exposed to a current, orienting and swimming according to the flow field. Consequently, flow-induced transport could be substantially different for the population members and any aggregating signal they secrete. With the aim of investigating how flows and rheotaxis responses impact on an aquatic population’s ability to form and maintain an aggregated profile, we develop and analyse a mathematical model that incorporates these factors. Through a systematic analysis into the effect of introducing rheotactic behaviour under various forms of environmental flow, we demonstrate that each of flow and rheotaxis can act beneficially or detrimentally on the ability to form and maintain a cluster. Synthesising these findings, we test a hypothesis that density-dependent rheotaxis may be optimal for group formation and maintenance, in which individuals increase their rheotactic effort as they approach an aggregated state.

## 1 Introduction

Living in an aquatic environment can expose an organism to strong and turbulent flows and it is natural to suppose means have evolved to exploit or counter them [1]. One known mechanism is *rheotaxis*, a response in which an individual reorients its body axis and swims with respect to the flow velocity field [2, 3, 4]. Positive rheotaxis indicates orientation and swimming against the current, which could allow an organism to hold its position, while negative rheotaxis implies swimming downstream, which could allow an organism to exploit the current for faster motion. Instances of rheotaxis have been recorded in a wide range of organisms, both at the level of single cells and large multicellular organisms. For example, rheotaxis has been observed for bacteria [5, 6], protozoa [2, 7] and mammalian sperm cells [4, 8], where in the latter rheotaxis may play a guidance role that orients sperm towards the egg. Rheotaxis has been widely studied for fish [3, 9, 10, 11, 12, 13, 14, 15], where it appears at scales ranging from larval zebrafish [13, 15] to whale sharks [16]. More generally, observations of rheotactic behaviour have been recorded for planktonic species [17], microscopic worms [18, 19], krill [20], jellyfish [21], salamanders [22], and turtles [23]. Much of this large literature, however, has focussed on individual-level rheotaxis, with relatively few studies exploring how rheotaxis interacts with and impacts on the movements designed to coordinate the collective behaviour of a population.

The temporary concentration of a normally dispersed population at some location can be an essential stage in the life cycle of a species [24, 25]. Benefits of aggregating range from efficient migration to productive hunting and feeding, or from protection against predation to reproduction. Many aggregations form routinely, as in the gathering of an established population at historical breeding grounds, but others appear unpredictably and appear to be driven by an innate self-organising process. One mechanism known to allow self-organisation of a population involves chemosensitive responses. Chemical communication between organisms is a near ubiquitous phenomenon, dictating numerous critical behaviours in aquatic populations [26]: both single cells and multi-cellular organisms are capable of detecting and responding to chemicals or odours, chemicals can be transported long distances from the signaller to receiver, and function in dark or noisy environments. Chemically-mediated self-organisation can occur through members of the population releasing an aggregating pheromone that attracts conspecifics, with evidence for this mechanism found in a range of unicellular [27, 28] and multicellular[29, 30, 31, 32, 33, 34, 35] terrestrial and aquatic organisms. As one example, the crown-of-thorn starfish (COTS, *Acanthaster planci*) releases water-borne factors that act to attract neighbours, subsequently generating an aggregate that (perhaps) initiates synchronised mass spawning [36]. Determining when, how and why such aggregates form is crucial for understanding the dynamics of an ecosystem; in the context of COTS, this specialised coral predator is capable of population outbreaks that subsequently decimate local reefs [37].

The reinforcing loop in which a population produces its own attractant will intuitively lead to aggregation of a population. Nevertheless, certain conditions must still be met before this positive feedback overcomes any negating processes that lead to dispersion, such as decay/loss of the aggregating signal. First, the population must be present in sufficient numbers for enough attractant to be produced. Second, population members must be sufficiently sensitive to allow them to detect and move towards an emerging source of attractant. For populations that reside in large domains, such as small organisms in a river or ocean environment, it is far from certain that these requirements will be satisfied when the population is scattered. The respective contributions of flow and rheotaxis add further uncertainty: while flow transports both the population and its attractant, rheotaxis counters, but only on the population. With their potential capacity to dramatically and distinctly alter the distribution of both the population and its attractant, it is therefore ambiguous whether flow and rheotaxis act beneficially or detrimentally on group formation and maintenance.

Motivated by this, we use modelling to systematically explore the extent to which flow and rheotaxis impact on aggregation dynamics in a population, specifically its capacity to (i) form, and (ii) maintain a clustered state. In the next section we describe the model, an adaptation of the well known Keller-Segel model for chemotaxis. The analysis starts with a simplistic uniform flow scenario, addressing the conditions under which a population can form aggregates in the absence and presence of rheotactic behaviour. We subsequently address post-aggregation dynamics, determining the tendency of clusters to unify and the level of rheotaxis required for a cluster to hold position. Exploring variable flow environments, we determine whether “favourable” flows can allow clusters to form under conditions where they would not usually do so, or whether “unfavourable” flows can destroy previously formed clusters. Noting the capacity of rheotaxis to maintain cluster integrity, we finally explore whether a density-dependent rheotactic responses can balance the positive and negative elements of each of rheotaxis and flow. We conclude with a brief discussion of the key results.

## 2 Model

We base our modelling on the well-known Keller-Segel system [38], a standard reaction-diffusion-advection model that has been adapted and applied to a broad spectrum of chemotactic aggregation processes (see [39] for a review). In particular, we extend it to incorporate flow and rheotaxis, see Fig. 1. Specifically,

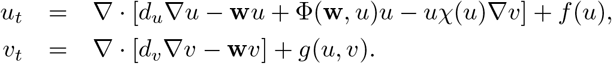

**Figure 1:**
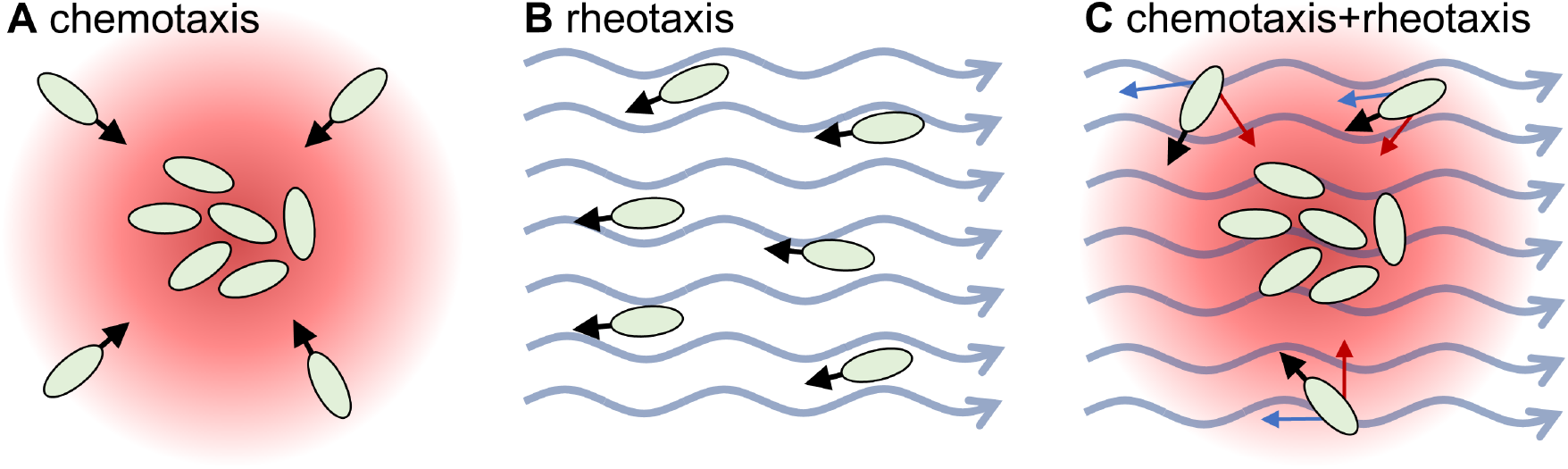
Schematic illustrating the principle components of the model. **A** Under chemotaxis alone, organisms are directed towards high concentrations of a secreted aggregating cue. **B** Under positive rheotaxis, organisms are transported by the flow, but also orient and swim against the current. **C** Under both rheotaxis and chemotaxis, the organisms must balance their overall movements according to both the attractant distribution and the flow field.

Here, *u*(**x**, *t*) and *v*(**x**, *t*) denote the population density and chemical attractant, respectively, defined at position **x** ∈ Ω ⊂ ℝ^*n*^ and time *t* ∈ [0, *T*]. For simplicity our study will be to restricted to simple spatial domains Ω: either a one-dimensional interval of length *L_x_* or a two-dimensional rectangular region of dimensions *L_x_ × L_y_*. *d_u_* and *d_v_* denote the population and chemoattractant diffusion coefficients, respectively. The chemotactic sensitivity *χ*(*u*) measures the strength of the chemotactic response and here we set *χ*(*u*) = *α* (1 − *u/k*_1_), where *α* is the chemotactic coefficient and density-limitation has been included to stop high densities (> *k*_1_) from forming: *k*_1_ represents the packing density [40]. We choose *f* (*u*) = *ru* (1 − *u/k*_2_) and *g*(*u, v*) = *βu − γv*, i.e. logistic growth of the population and linear secretion/decay of the attractant. These assumptions form a relatively standard set for describing biological aggregation phenomena, e.g. [39].

Flow is included as a prescribed vector field, **w**(**x**, *t*), that transports both the population and its attractant. Rheotaxis is modelled as a further directed movement of the population, encoded in a function Φ(**w**, *u*) that depends on the flow field and population density. Here we will consider two forms for this function:

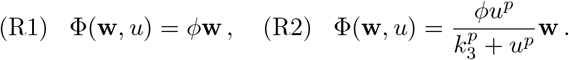

For either choice, rheotactic movement is upstream (downstream) for rheotactic coefficient *ϕ* > 0 (*ϕ* < 0). (R1) describes a simple linear relationship between rheotaxis effort and the magnitude flow, while (R2) extends this to incorporate a rheotaxis response that also increases with the (local) population density; in (R2) *k*_3_ defines the point at which rheotaxis behaviour is engaged, with *p* the Hill coefficient. Rheotaxis is distinctly interpreted for free-swimming or bottom-dwelling organisms, where (assuming the linear choice, R1) for the former *ϕ* = 1 implies a compensating response in which upstream swimming balances downstream flow, while for the latter *ϕ* = 1 implies fixing to the floor. *ϕ* > 0 denotes positive rheotaxis; *ϕ* = 1 indicates compensating rheotaxis; regimes 0 < *ϕ* < 1 and *ϕ* > 1 describe undercompensating and overcompensating responses, respectively. At the boundaries of Ω we will impose periodic conditions which, while somewhat artificial, limit the potential for boundary accumulations while preserving population mass. Note that the domain itself will be chosen to be large with respect to the characteristic length scale of clusters. Initially we set *u*(**x**, 0) = *u*_0_(**x**) and *v*(**x**, 0) = *v*_0_(**x**), further details of which are provided below. By rescaling (*u* = *k*_1_*u*\*, *v* = *βk*_1_*v*\*/*γ*, 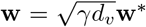, *t* = *t*\*/*γ*, 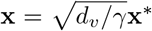, *δ* = *d_u_/d_v_*, *α*\* = (*αβk*_1_)/(*d_v_γ*), *U* = *k*_2_/*k*_1_, *κ* = *k*_3_/*k*_1_ and *ρ* = *r/γ*) and dropping stars, we arrive at the non-dimensional system of equations

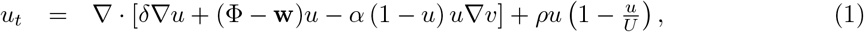

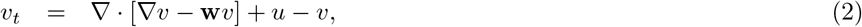

with either (R1) Φ = *ϕ***w** or (R2) 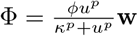.

Equations (1–2) have a positive uniform steady state (USS) at (*U, U*). Note that when population growth is negligible (*ρ* = 0) and lossless boundary conditions are assumed, the USS will be instead determined by the average initial population density, which we also denote by *U*. *U* therefore represents the mean dispersed density and we will define a population as *clustered* at position **x** and time *t* if *u*(*x, t*) ≥ 4*U*, a value compatible with definitions for a marine spawning aggregation [41]. According to the case under consideration, initial distributions *u*_0_(**x**) and *v*_0_(**x**) will be formulated in either a dispersed or clustered state. Dispersed initial conditions imply a population that is initially nonclustered and quasi-uniformly distributed about the steady state *U*. Clustered initial conditions describe a population for which there are initially one or more regions where *u*(**x**, 0) > 4*U*.

The rest of the paper focusses on the dynamics of (1–2), in particular under variation of:

1. flow velocity field **w**(**x**, *t*), parametrised by *maximum flow speed ω*;
2. chemotactic coefficient *α*, a measure of the strength of the aggregation mechanism;
3. rheotactic coefficient *ϕ*, measuring the rheotactic effort;
4. dispersed density *U*, measuring the overall size of the population.

Flow fields will be described as uniform (nonuniform) if they are independent of (depend on) **x** and constant (nonconstant) if independent of (depend on) *t*. Uniform-constant flows are primarily selected for analytical convenience, nevertheless they may approximate certain environments over shorter timescales (such as a steadily flowing stream) or laboratory experiments of rheotactic behaviour, e.g. [42]. For computational convenience we do not explicitly solve a Navier-Stokes equation to generate the flow, rather we prescribe **w**(**x**, *t*) functionally or via a dataset (see Appendix).

## 3 Results

### 3.1 Autoaggregation in uniform flows

For simplicity we begin with uniform/constant flow in a quasi one-dimensional setting (e.g. the length of a stream), thus **w** = *ω* ≥ 0 (left to right flow). Excluding rheotaxis (*ϕ* = 0), standard linear stability analysis (LSA, see Appendix) predicts autoaggregation of the dispersed population into clusters if the following simple threshold is met:

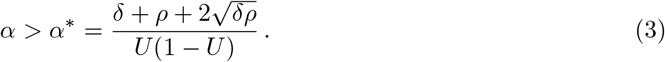

See Fig. 2**A** for a representative parameter space. Given condition (3) a dispersed population organises into clusters separated by a characteristic wavelength, as highlighted by representative simulations in Fig. 2**B-C**. As noted in the introduction, this well-known instability arises from positive chemotactic feedback: autoaggregation is only possible if (i) the population lies above a critical density, (ii) the chemotactic response is sufficiently strong, (iii) the population produces sufficient attractant. Regimes *α > α*\* and *α < α*\* are referred to here as strong and weak chemotaxis scenarios, respectively, according to whether chemical communication is strong enough to induce clustering. Note that (uniform) flow does not alter condition (3), but does result in downstream movement of clusters (compare Fig. 2**B** with Fig. 2**C**).

**Figure 2:**
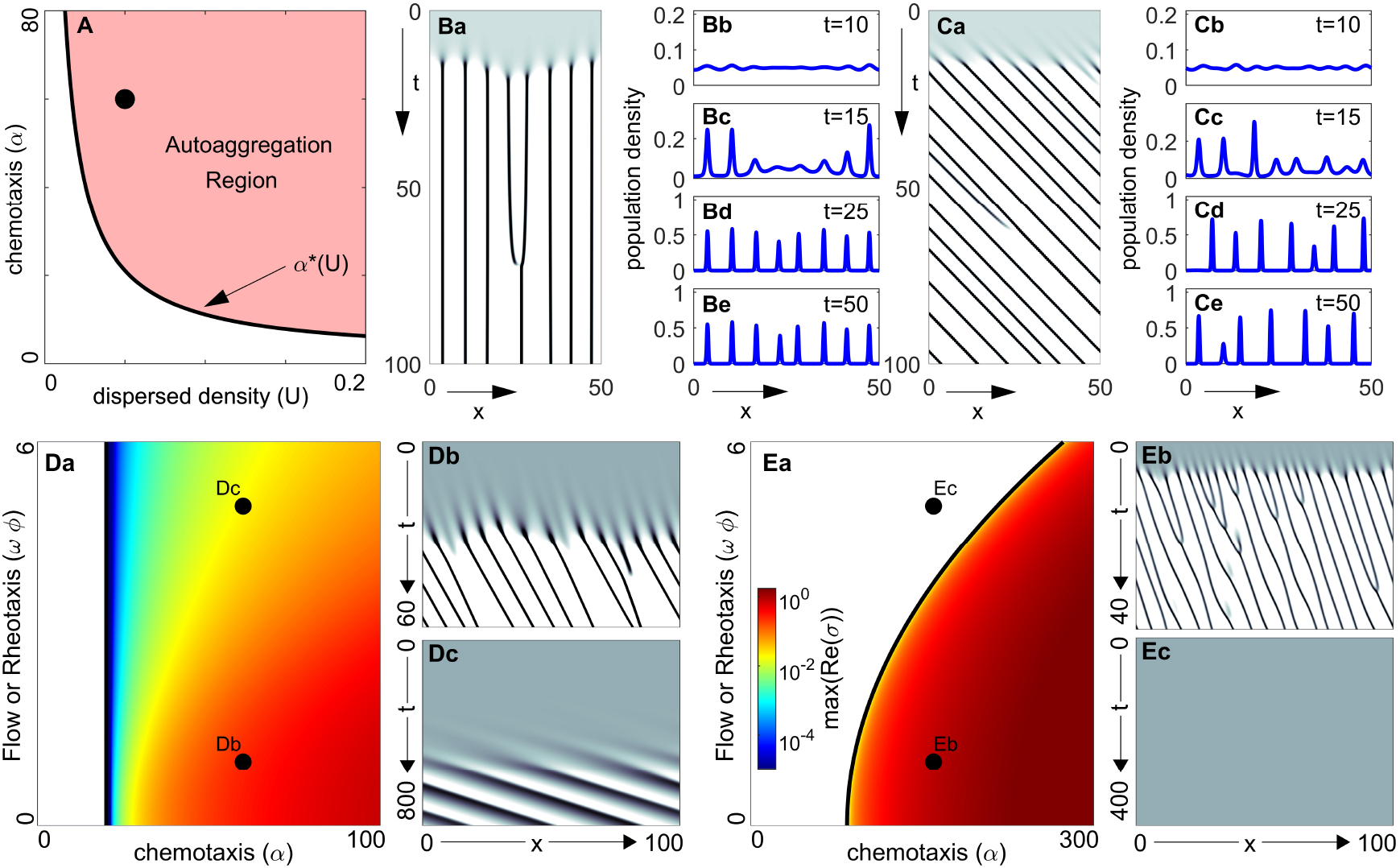
Aggregation in a uniform flow. **A** Autoaggregation parameter space (*α* - *U*), with parameters for plots in **B-C** indicated by dot. Note that here we assume negligible population growth (*ρ* = 0). **B** Autoaggregation in a still environment, *ω* = 0. Population density *u* represented as **a** space-time map (white *u*(*x, t*) = 0, black *u*(*x, t*) ≥ 4*U*) and snapshots in **b-e**. **C** As **B**, but under constant flow (*ω* = 1). **D** Autoaggregation parameter space (*α* - *ϕω*, coloured region) under rheotaxis and flow (zero population growth, *ρ* = 0). Colour (see inset of **Ea**) indicates the cluster growth rate predicted by LSA (deep red = faster growth). Space-time density maps for marked points: **Db** *α* = 60, *ω* = 1, *ϕ* = 1; **Dc** *α* = 60, *ω* = 4, *ϕ* = 1. **E** As **D** but including population growth (*ρ* = 1). **Eb** *α* = 150, *ω* = 1, *ϕ* = 1; **Ec** *α* = 150, *ω* = 4, *ϕ* = 1. For all simulations we have set initial densities as dispersed (*u*_0_(*x*) = *U, v*_0_(*x*) = *U* + *E*(*x*), for small random perturbation *E*(*x*)). Nonspecified parameters are *U* = 0.05 and *δ* = 1. Details of the LSA are provided in the Appendix.

### 3.2 Rheotaxis subdues clustering

When rheotaxis (*ϕ* ≠ 0) is included, condition (3) remains necessary for autoaggregation to occur (see Appendix). This suggests that there is no expansion of the parameter regime in which clustering can occur (weak chemotaxis remains insufficient to induce aggregation of a dispersed population). When we exclude population growth (i.e. *ρ* = 0) condition (3) turns out also to be a sufficient condition, given the assumption of an effectively infinite domain (such as a long stream): we can observe this via the independence between the size of the autoaggregation parameter space and the size of *ϕ* in Fig. 2**Da**. While this suggests that adding rheotaxis does not alter the fundamental ability of a dispersed population to organise into clusters, rheotaxis does in fact suppress autoaggregation through dramatically restricting the cluster growth rate (see Appendix for its definition): observe the order of magnitude time difference before dense clusters are formed in Fig. 2**Db** (low flow/rheotaxis combination) and Fig. 2**Dc** (high flow/rheotaxis combination). This growth-retarding consequence of rheotaxis could prevent significant accumulation of a population within biologically or ecologically relevant timescales.

The inclusion of population growth (*ρ* > 0) leads to an even more obvious impact from rheotaxis, with clustering now abolished above a critical rheotaxis/flow combination even under strong chemotaxis. Fig. 2**Ea** plots a representative autoaggregation parameter space when *ρ* > 0, where the strong dependency on the rheotactic effort *ϕ* is observed. Selecting parameters from a strong chemotaxis regime, simulations indicate that a larger level of rheotaxis can abolish clustering, compare Fig. 2**Eb** and Fig. 2**Ec**. Summarising, engaging in (simple) rheotaxis within an effectively uniform flow appears to be counterproductive for forming aggregates through chemotaxis.

### 3.3 Rheotaxis accelerates unification

We next explore the post-aggregation behaviour, i.e. how flow and rheotaxis alter the dynamics of previously formed clusters. Note that herein we restrict to negligible population growth (*ρ* = 0), effectively assuming that growth is negligible on the timescale of aggregation behaviour. In the absence of rheotaxis, chemotaxis autoaggregation processes are often characterised by a series of *unifying* events, where neighbouring clusters attract each other and merge, leading over time to fewer, but larger aggregates (for example, see [43]); this could be perceived as beneficial for a population aiming to form very large clusters. Unification time (which we define as the time taken to evolve to a single cluster) under chemotaxis alone, though, may be unrealistically long, due to the reliance on diffusion of the attractant through the inter-aggregate space; in the representative simulation (Fig. 3**Aa**) we observe that none of the clusters that initially form have merged by the end of the simulation, an order of magnitude longer than their formation time. In contrast, when rheotaxis is incorporated we find that unification is emphatically accelerated, with multiple merging processes resulting in just one or two clusters over the same timespan (Fig. 3**Ab-c**). Averaged across multiple simulations, we observe a multiple order of magnitude reduction in the unification time as rheotaxis is increased, Fig. 3**B**. Summarising, the addition of rheotaxis appears to have a dramatic capacity on the post-aggregation unification of clusters.

**Figure 3:**
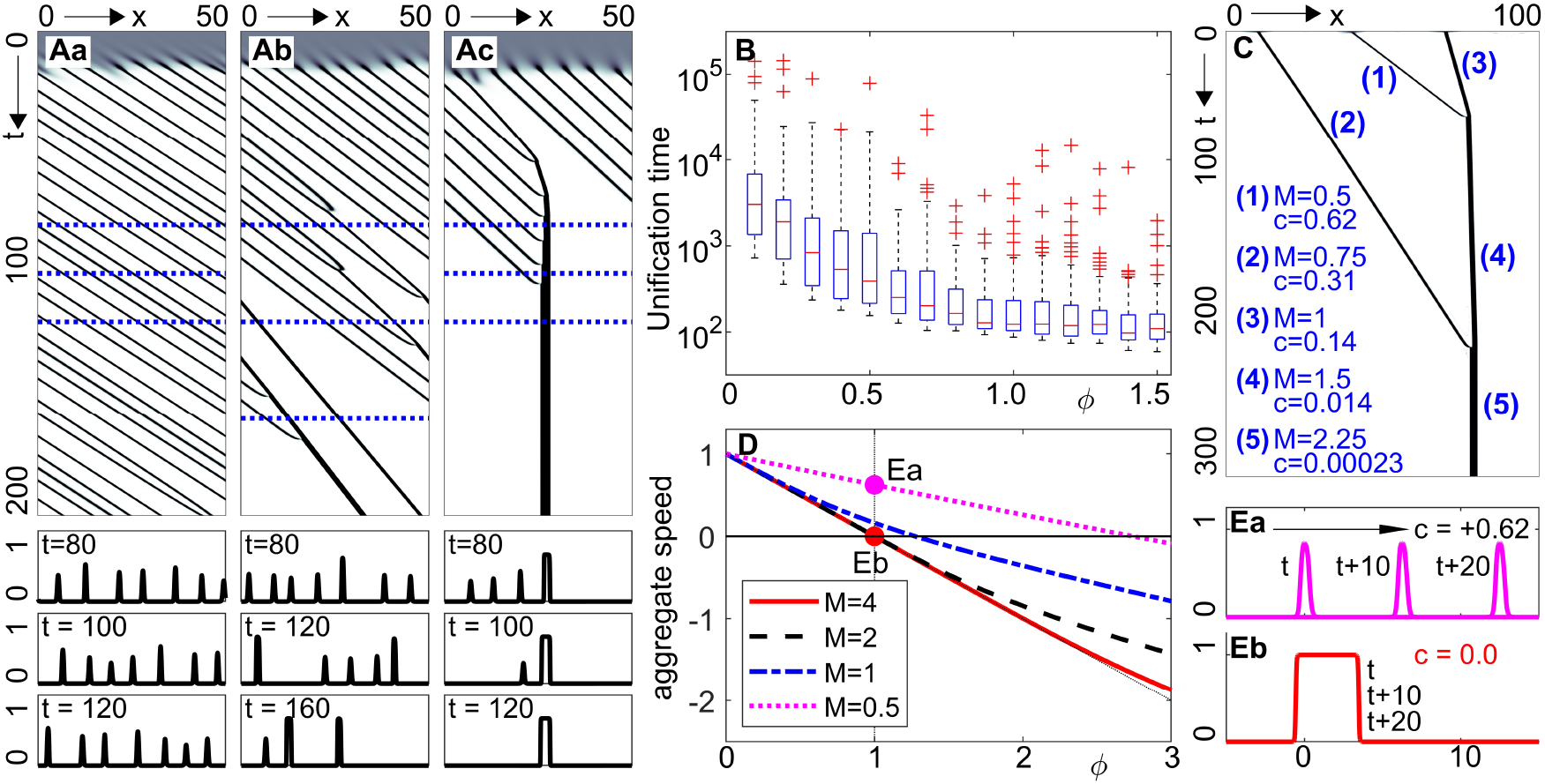
**A-B** Accelerated unification through the action of rheotaxis. **A** Space-time population density maps showing cluster evolution under a constant flow (*ω* = 1) and **a** negligible (*ϕ* = 0), **b** under-compensating (*ϕ* = 0.5), **c** compensating rheotaxis (*ϕ* = 1). **B** Boxplot showing the mean time to reach a unified cluster (averaged across 100 randomised initial data). For **A-B** initial distributions are as in Fig. 2. **C** Evolving density for a population initialised as three separated Gaussian shaped clusters of masses *m* = 0.5, 0.75 and 1, centred at *x* = 40, 10 and 70 respectively. **D** Computed cluster speed, *c*, as a plot of rheotaxis strength, *ϕ*, for isolated clusters of mass *m*; dotted vertical line indicates compensating rheotaxis and horizontal solid line indicates holding position. **E** Travelling pulse profile for a cluster of mass **a** *m* = 0.5 and **b** *m* = 4 under compensating rheotaxis. Density *u*(*x, t*) shown at *t* = *t*\*, *t*\* + 10, *t*\* + 20, with arrow indicating movement direction. In **A-E** *ρ* = 0, *δ* = 1 and *α* = 80.

This accelerated unification stems from equivalent rheotaxis (same effort for each member, regardless of group status) acting distinctly on different sized clusters. Pertinently, a large cluster will hold position more effectively than a small cluster and, consequently, the latter drifts into and merges with the former. We demonstrate this phenomenon in Fig. 3**C**, initially distributing the population into spatially separated clusters of distinct masses. Under compensating rheotaxis (*ϕ* = 1), the smaller-sized cluster is significantly transported by the flow, the medium-sized less so and the large cluster almost holds position. Over time, clusters merge into a single large group that remains quasi-motionless. To investigate this further, we numerically evaluate cluster wavespeed (*c*) as a function of rheotaxis strength and cluster mass (*m*), where *c* = 0 indicates a cluster holding position and *c* > 0 (*c* < 0) implies transport downstream (upstream). Results (Fig. 3**D**) consistently yield *c*(*ϕ, m*_1_) > *c*(*ϕ, m*_2_) for *m*_1_ < *m*_2_. A corollary of this is that it is easier for an individual to be in a larger cluster to maintain position: small clusters may require overcompensating responses (i.e. *ϕ* > 1) for the cluster to hold position. An explicit simulation is shown in Fig.3**E**, where for the same rheotaxis the small mass shifts downstream while the large mass holds position. Summarising, one can perceive a potential benefit of forming a larger group when it comes to maintaining position against a flow.

### 3.4 Nonuniform flows facilitate clustering

As a controlled nonuniform flow we consider a uniform flow which becomes temporarily interrupted by a vortex structure, see Fig. 4**A**. Excluding chemotaxis (*α* = 0) and rheotaxis (*ϕ* = 0), this flow generates a population that remains, in essence, dispersed: while introduction of the vortex flow temporarily structures the population through its moderate accumulation inside sluggish regions, the population density remains significantly below the threshold used to define clustering (Fig. 4**B**). When we combine this flow environment with even just weak chemotaxis, however, we can observe clustering, Fig. 4**C**: the moderate flow-induced accumulation becomes amplified through positive chemotaxis feedback and the population rounds into a cluster. Moreover, once formed this cluster is able to persist even when the vortex has been removed and flow returns to uniform.

**Figure 4:**
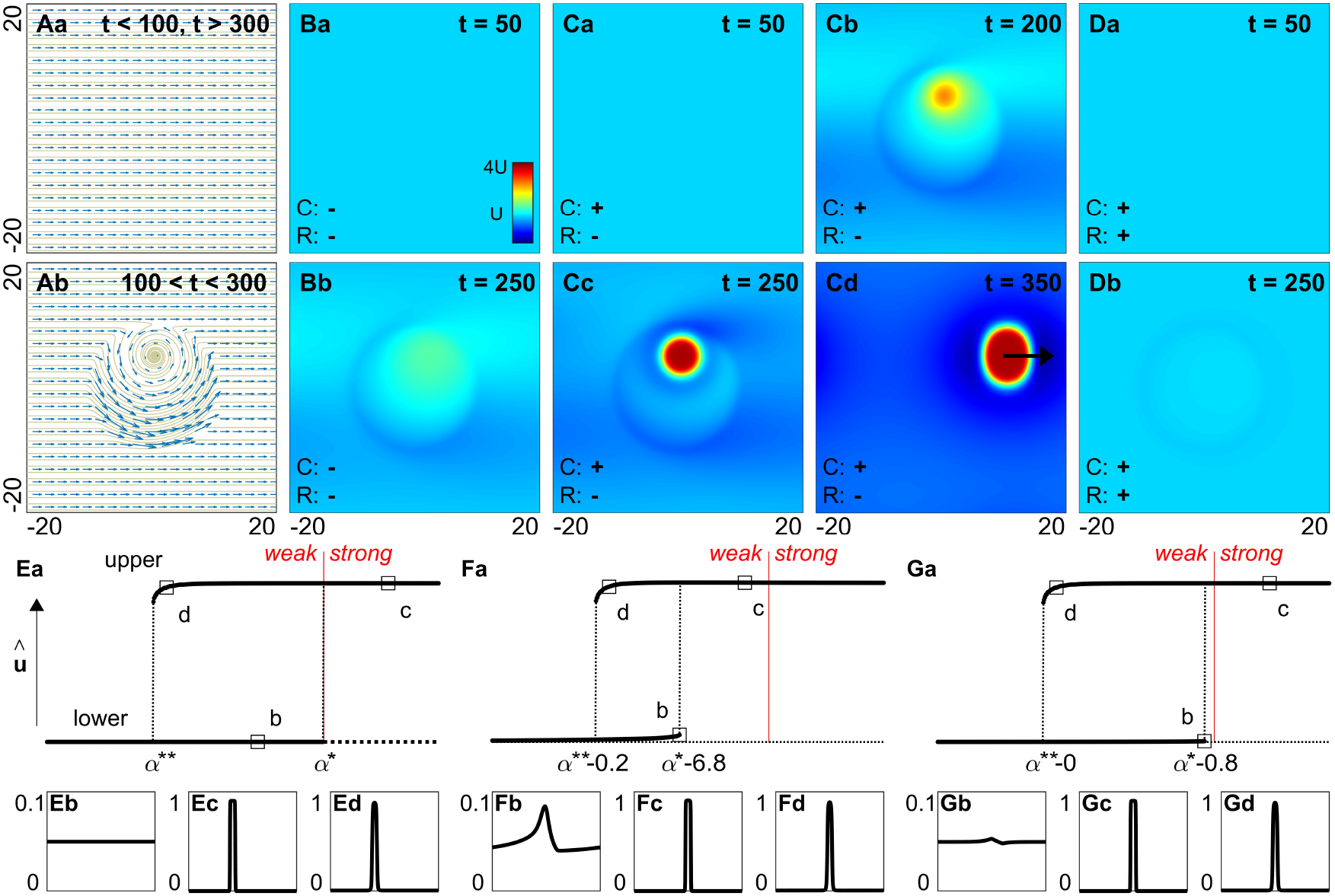
Dynamics under nonuniform flow. **A** The flow field, **w**. **B-E** density *u* (colourscale in **Ba**). **B** Excluding chemotaxis and rheotaxis (*α* = *ϕ* = 0), the population stays unclustered, though nonuniform flow leads to moderate accumulations. **C** Under weak chemotaxis and no rheotaxis (*α* = 10, *ϕ* = 0), flow-induced accumulations form into clusters that are sustained following removal of the vortex flow. **D** Addition of rheotaxis (*α* = 10, *ϕ* = 1) suppresses the clusters that formed in **C**. In **A-D** the population is initially dispersed, *u*_0_(**x**) = *U, v*_0_(**x**) = *U* + *ϵ*(**x**). **E-G** Bifurcation diagrams of numerically-determined steady states, *u_s_*(*x*), represented via 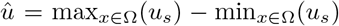. Bifurcation parameter is *α*. Solid branches indicate numerically stable steady states, dotted lines indicate unstable steady states. Representative steady states for the locations indicated by a square shown in below panels. **E** Uniform flow, no rheotaxis (*φ* = 0). **F** Uniform flow, interrupted by a region of slower flow, no rheotaxis (*ϕ* = 0). **G** As **F**, but with rheotaxis (*ϕ* = 1). Simulations in **A-D** use Ω = [−20, 20] × [−20, 20] and *w*(*x, y, t*) = (1, 0)+0.005(−*x*−10*y,* 10*x−y*)[tanh(*t* − 100)−tanh(*t*−300)][1 − tanh(0.1*x*^2^+0.1*y*^2^−10)]. Simulations in **E-G** use Ω = [−50, 50] and *w*(*x*) = 1.0 + *ε*(1.0 − tanh(*x*) + tanh(*x* − 10)), where **E** *ε* = 0, **F-G** *ε* = 0.01. In all simulations, *U* = 0.05, *δ* = 1 and *ρ* = 0.

In other words “advantageous” nonuniform flows permit clusters to form and persist in a scenario where quasi-uniform (or negligible) flow would not. We attribute this phenomenon to hysteresis behaviour in the chemotaxis autoaggregation mechanism. To test this we employ a numerical bifurcation analysis (following the method in[43]) within an idealised one-dimensional simulation, specifically where a uniform left to right flow is interrupted by a localised region of slow flow. Under uniform flow the bifurcation structure of the autoaggregation model can be summarised by two principal solution branches: a lower branch representing the USS and an upper branch representing a clustered population, see Fig. 4**E**. Passing from weak to strong chemotaxis (at *α* = *α*\*) destabilises the USS and chemotactic feedback powers accumulation into a cluster (as predicted by the LSA). The clustered branch is stable either direction about *α*\*, a hysteresis phenomenon which implies that once a cluster has formed it maintains structure even if chemotaxis drops back to a weak level, down to a lower threshold at *α*\*\*.

The addition of nonuniform flow to this induces a “wrinkling” of the uniform distribution, subsequently raising the local density above the threshold needed for chemotaxis to initiate autoaggregation. As a consequence, the critical threshold at which the lower branch becomes unstable decreases, Fig. 4**F**, and emergence of clusters from a dispersed state now occurs for weak chemotaxis. Once settled on the upper branch, the hysteresis phenomenon means that the cluster will remain stable even if the nonuniform flow is removed.

How does rheotaxis impact on this? Notably, rheotactic behaviour counteracts flow-induced accumulations and therefore drives the critical threshold back towards the weak/strong chemotaxis boundary, even when the flow is highly nonuniform, Fig. 4**G**. Consequently, while nonuniform flows encourage dense clustering in populations that do not exhibit rheotaxis, enabling rheotaxis suppresses this potentially beneficial outcome. This is confirmed numerically in Fig. 4**D**, where the addition of compensating rheotaxis prevents the formation of the cluster observed in Fig. 4**C**.

### 3.5 Rheotaxis prevents cluster disintegration

The above highlighted a potential benefit of variable flow (nudging local densities beyond a critical threshold required for autoaggregation) that was negated when individuals engaged in rheotaxis. Here we show the reverse: a destructive consequence of variable flow neutralised by rheotaxis. We impose a quasi-realistic nonuniform/nonconstant flowfield **w**(**x**, *t*) (see Appendix for details), parametrised by a magnitude parameter (*ω*) that indicates the maximum flow speed experienced. We initially arrange a population in a clustered form and choose parameters from the weak chemotaxis regime; note that the weak chemotaxis is sufficient to hold the cluster together in the absence of flow/rheotaxis, through the above described hysteresis.

We explore cluster evolution, measuring two time-dependent quantities: the proportion remaining clustered and the proportion remaining at the initial cluster zone (definitions in Fig. 5 caption). Under a weak flow field, Fig. 5**A**, cohesion of the cluster is maintained despite some distortion and drift away from the initial clustering zone. For moderate to strong flow fields, Fig. 5**BC**, we observe more substantial drift, along with splintering of the cluster. Here the varying flow field starts to pull the cluster in different directions, smaller groups are peeled from the main group and they either reattach, remain separated, or disintegrate and collapse. We observe rapid decline in the capacity to remain localised and, for stronger flows, a decrease in the proportion that remain clustered. Note that similar results are observed under strong chemotaxis scenarios (data not shown).

**Figure 5:**
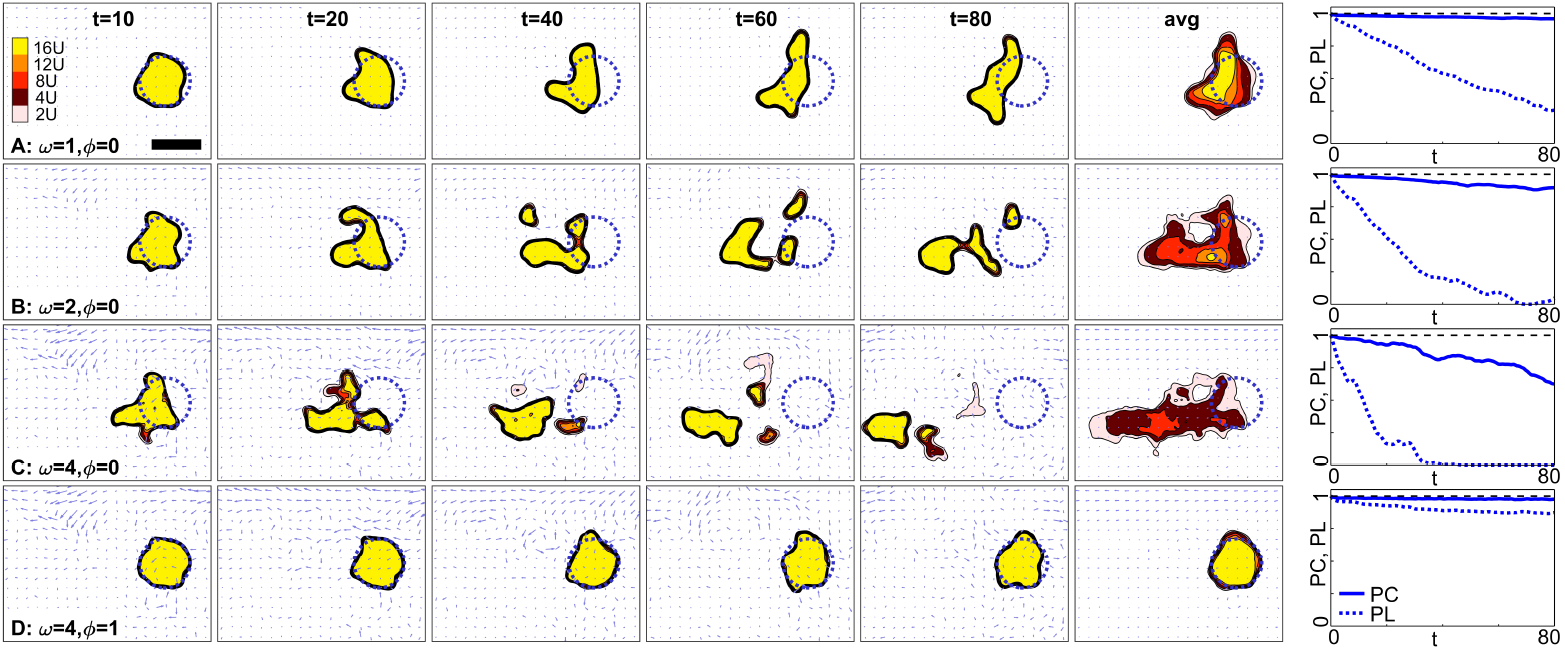
Nonuniform, nonconstant flows disintegrate clusters, but they can be stabilized by rheotaxis. **A-C** Nonuniform, nonconstant flow and zero rheotaxis (*ϕ* = 0). Each frame shows the instantaneous flowfield (arrows), population density (colourscale) and initial cluster location (dotted blue circle) under: **A** weak flow, *ω* = 1; **B** moderate flow, *ω* = 2; **C** strong flow, *ω* = 4. **D** Nonuniform, nonconstant flow and compensating rheotaxis (*ϕ* = 1), under strong flow, *ω* = 4. The final column plots two measures: the proportion clustered (PC) and the proportion localised (PL), both normalised against their values at *t* = 0. Solid bar in top left represents a length scale of 10. **avg** shows densities averaged across time from *t* = 0 to 80. Initial circular cluster centred on (0, 0) and given by *u*_0_(*x, y*) = *v*_0_(*x, y*) = 0.5(1 − tanh(10(*x*^2^ + *y*^2^ − 25))) and we set *ρ* = 0, *δ* = 1, *α* = 15 (weak chemotaxis regime). *PC*(*t*) = ∫_Ω_ *H*(*u*(**x**, *t*) − 4*U*)*u*(**x**, *t*)*d***x**, where Ω is the full spatial region and *H*(·) denotes the Heaviside function. 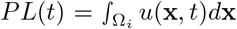, where Ω_*i*_ denotes the region inside the dotted blue line. Flowfield uses the HYCOM dataset described in methods.

Rheotaxis, of course, can act as a counter to flow and we examine the extent to which a compensating rheotaxis response (*ϕ* = 1) impacts on these dynamics. Notably, the cluster-destabilising effects of flow are negated when the population performs rheotaxis: in such scenarios the aggregate remains clustered and more or less localised to the initial cluster zone, see Fig. 5**D**.

### 3.6 Optimised clustering via density-dependent rheotaxis

As we have shown, flow and rheotaxis can be positive and negative when it comes to forming and maintaining clusters. Benefits of flow were found in forming clusters, with repercussions on cluster maintenance; for rheotaxis it was the reverse, limiting autoaggregation but unifying and stabilising clustered populations. This leads us to hypothesise that density-dependent rheotaxis may optimise aggregation formation and maintenance, specifically a response in which individuals increase their rheotactic response according to population density.

To test this we consider the density-dependent rheotaxis form *φ* = *u^p^*/(*κ^p^* + *u^p^*), modelling low to negligible rheotaxis at unclustered densities and approaching compensating rheotaxis for clustered populations. Note that *κ* = 0 indicates constant compensating rheotaxis and *κ* = ∞ indicates negligible rheotaxis. We initialise the population as dispersed, select from the weak chemotaxis regime and choose the strong flow setting used in Fig. 5.

Under constant compensating rheotaxis, Fig. 6**A**, any flow-induced inhomogeneities are suppressed and the population remains unclustered: the density remains below instability inducing thresholds. Under negligible rheotaxis, on the other hand, flow-induced inhomogeneities are amplified into clusters through chemical communication, yet flow subsequently distorts and transports these through space, Fig. 6 **D**. The introduction of density-dependent rheotaxis responses can permit both the formation of clusters and their subsequent maintenance, Fig. 6 **B-C**. In their dispersed state, individuals barely engage in rheotaxis and subsequently the population benefits from flow-induced local accumulations that induce clustering. Once clustered, compensating rheotaxis is engaged that leads to an essentially stable cluster that maintains its position in space.

**Figure 6:**
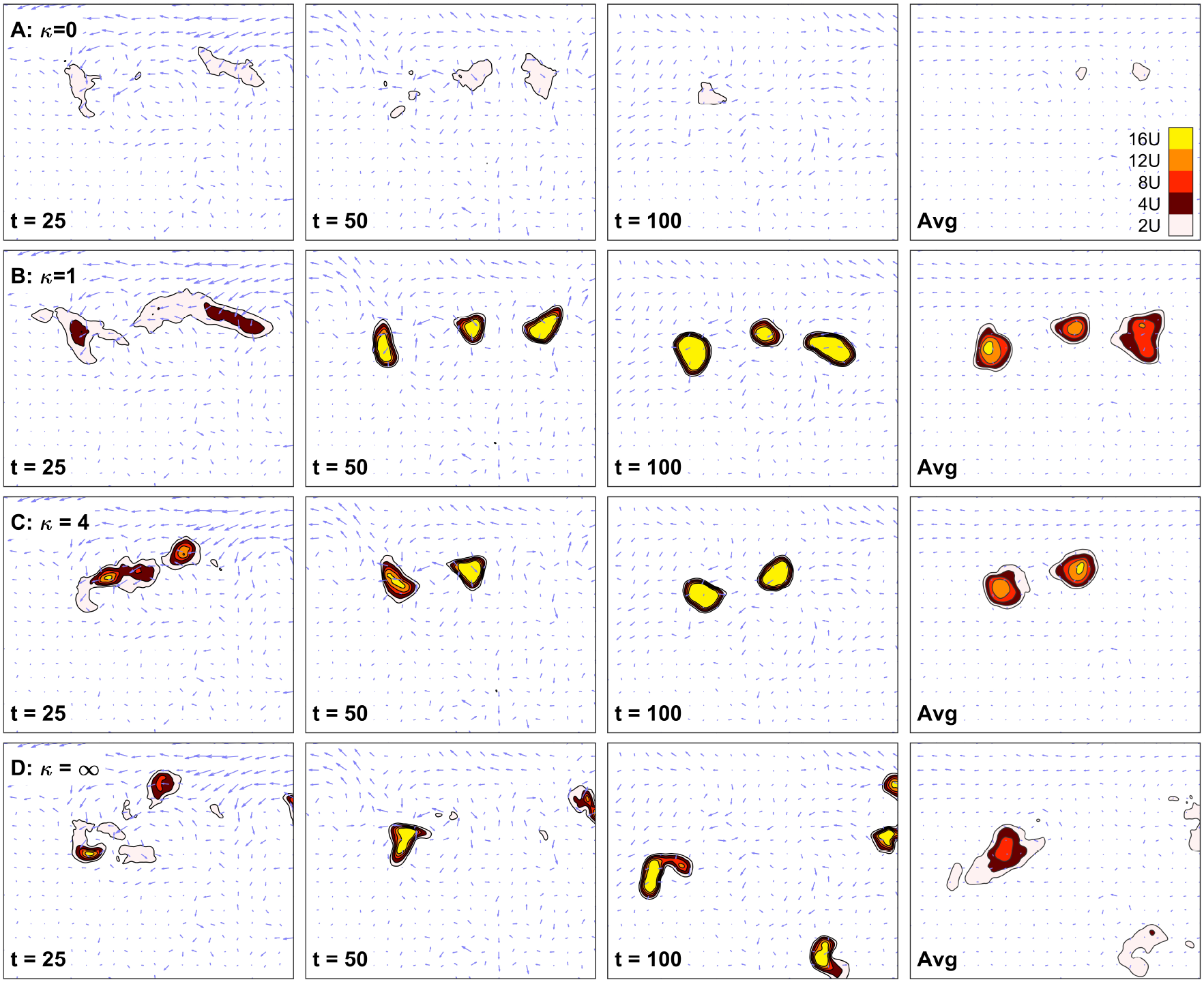
Dynamics of an initially dispersed populations exhibiting density-dependent rheotaxis. Each panel shows the instantaneous flow field (arrows) and population density *u*, (colourscale: top right). Strong flow from Fig. 5. Rheotaxis coefficient *φ* = *u^p^*/(*k^p^* + *u^p^*), for: **A** *κ* = 0 (constant compensating rheotaxis); **B** *κ* = 1; **C** *κ* = 4; **D** *κ* = ∞ (zero rheotaxis). Initial distributions *u*_0_(*x*) = *U, v*_0_(*x*) = *U* + *ϵ*(*x*) and we use *p* = 2, *ρ* = 0, *δ* = 1, *U* = 0.05, *α* = 15 (weak chemotaxis regime). Flowfield uses the HYCOM dataset described in methods.

## 4 Discussion

Aggregating may be required at various stages of a population’s life cycle, yet when and how individuals find their way into groups can be difficult to deduce. Chemical signalling is an ancient and near-ubiquitous mode of communication [44], which can allow a population to group through communal secretion of an aggregation pheromone. A requirement is that the dispersed density is sufficiently high, so that enough attractant is produced for neighbours to move into and reinforce a developing cluster. Fully dispersed populations may lie below this threshold, raising the question of how a critical density is initially achieved. External flow provides a mechanism, bringing individuals to a number that seeds the aggregation. Similar observations have been observed in other models for animal grouping, for example [45].

On the other hand, flows can also fragment an aggregate, disadvantageous if the purpose of the cluster has not been achieved. We have shown that if a clustered population performs positive rheotaxis then the aggregate remains stable. Further, rheotaxis hastens cluster unification, beneficial if it is optimal to form the largest possible aggregate, e.g. mass spawning. Larger groups also require less rheotaxis to maintain position within a flow, implying energy expenditure benefits of being in a large group. Balanced against these advantages, though, is that rheotaxis slows cluster growth rate and suppresses the flow-induced seeding of aggregations. On the basis of these observations we have tested a hypothesis that density-dependent rheotaxis responses can optimise aggregate formation and maintenance. Under this conjecture, zero rheotaxis led to unstable clusters, constant rheotaxis entirely suppressed clustering, but density-dependent rheotaxis allowed persisting clusters to form from an initially dispersed state. Previous modelling studies have shown that improved rheotaxis performance may be a natural outcome of grouping, via close proximity and neighbour alignment reducing uncertainty [46].

A natural application is broadcast spawning, a common reproduction method used by marine organisms and involving the synchronised release of male and female gametes into the water column. Aggregation is logical, as higher densities will locally increase gamete concentration and fertilization rates [47]. Various studies have explored the impact of flow on broadcast spawning success, e.g. see [48], but at the level of gamete dynamics rather than population grouping. Natural extensions of our work here could include integrating it with equations for gamete dynamics, or exploring the impact of distinct chemical responses by distinct sexes [35]. Aggregating for spawning, though, comes at the obvious cost of conspicuousness to predators, illustrated by the tendency of fisheries to target fish spawning aggregations [49]. A reverse question to that posed in the present work, therefore, would be to address whether certain responses could facilitate rapid dispersal of a group once its purpose has been achieved.

Animals use various forms of communication to generate groups, and the focus on chemically-mediated aggregation here is perhaps particularly relevant to organisms with limited sensory systems, e.g. certain microorganisms and marine invertebrates. Fish, mammals *etc* can also rely on vision, sound and other cues, and a number of models have been constructed that describe the collective dynamics of such populations, e.g. [25]. Flow and rheotaxis are likely to play a substantial role on, say, fish shoals and a key question lies in whether rheotaxis and flow can similarly encourage shoal formation and maintenance. We have also taken a purposefully naive approach to fluid interactions, restricting fluid dynamics to a transport-only impacting role on the population and hence neglecting any feedback that results from the group on the flow field. More sophisticated coupled chemotaxis-flow systems have been developed [50, 51, 52], and these could be extended to account for behaviours such as rheotaxis. Nevertheless, we believe the present study acts as a starting point for an understanding the complexities between communication, orientation responses and flow on organism grouping.

## Acknowledgements

KJP thanks Thomas Hillen for comments on an early draft and acknowledges departmental funding through the “MIUR–Dipartimento di Eccellenza” programme.

## Appendix

## 4.1 Quasi-realistic flow fields

Functions for constant flows are stated in the relevant figure caption. For the quasi-natural flows used in Figs 5–6 we use a publicly available dataset (global HYCOM, a validated ocean forecasting model [53]). While HYCOM data is used for ocean scale dynamics, our modelling is nondimensional and we abstract it to nondimensional space and time scales, parametrised by a single parameter (*ω*) that denotes the maximum flow speed. Fundamentally, HYCOM generates natural flow patterns with typical features including semi-persistent currents and localised eddies, providing a plausible real world dataset. To limit boundary effects, this nonuniform, nonconstant flow data is “immersed” within a larger field of uniform and constant flow and simulations focus on dynamics near the central region, away from boundary-induced artefacts.

## 4.2 Linear stability analysis (LSA)

For simplicity we consider the non-dimensional system (1-2) on a 1D infinite line, uniform flow **w**(*x, t*) = *ω* and Φ = *ωϕ*. Standard LSA [54] about the USS at (*U, U*) yields stability matrix,

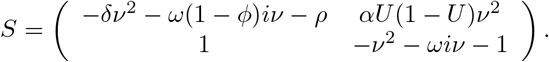

*ν* ≥ 0 is the spatial wavenumber (inversely related to the pattern wavelength). When *S* has (at least one) eigenvalues (*σ*) with positive real part for (at least one) valid *ν* then the USS is unstable and self-organisation/autoaggregation may occur; note that for the infinite line, all nonnegative real numbers are valid wavenumbers. We denote by max (Re(*σ*)) as the largest positive real part of eigenvalues across all valid wavenumbers, and note that instability therefore occurs when max (Re(*σ*)) > 0. We note further that the size of max (Re(*σ*)) can be viewed as a measure of the cluster growth rate, i.e. how quickly clusters will emerge from an almost uniform distribution. If *ϕ* = 0 then the condition for max (Re(*σ*)) > 0 is simply when (3) holds.

An explicit condition is difficult to obtain when *ϕ* ≠ 0 due to the presence of imaginary components. Stability is instead analysed through the neutral stability curves (NSCs, the curves of Re(*σ*) = 0, e.g. [55]). A bit of algebraic rearrangement determines NSCs as

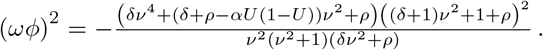

The right hand side (as a function of *ν*^2^) determines whether unstable wavenumbers exist at a given *ωϕ*, specifically according to whether NSCs bisect the positive quadrant of the |*ωϕ| − ν*^2^ plane. When population growth is negligible (*ρ* = 0) and the domain is infinite, a necessary condition for this to occur is simply that (3) holds, regardless of the size |*ωϕ*|, see Fig. 7**A**: given the infinite domain and condition (3), autoaggregation therefore remains possible under the inclusion of rheotaxis. However, it is noted that as the size of *ωϕ* increases, the range becomes increasingly restricted to small *ν*. Effectively, this restricts emerging clusters to those with long wavelengths and slow growth.

**Figure 7:**
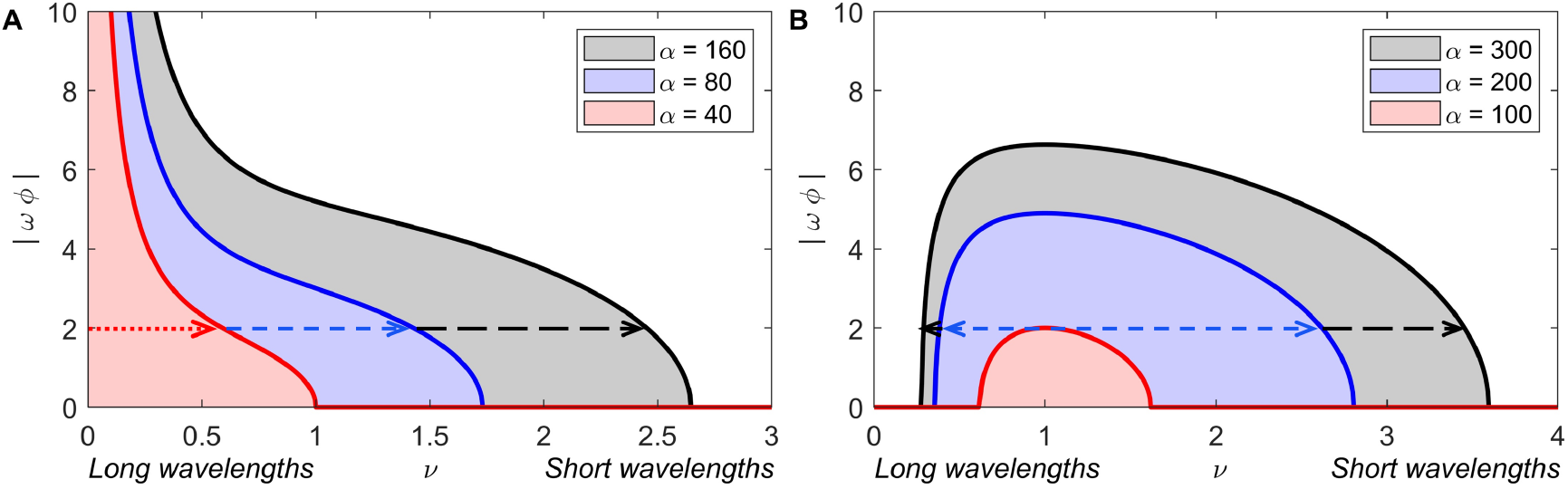
Neutral stability curves (NSC), indicating the range of unstable wavenumbers for particular (*ω*,*ϕ*) combinations: non-zero length ranges indicate the possibility of autoaggregation from the USS. **A** Zero population growth (*ρ* = 0), for three choices of *α*. **B** Logistic population growth (*ρ* = 1), for three choices of *α*. Under zero population growth a non-zero length range is found for all (*ω*,*ϕ*) combinations, although of restricting length as these parameters increase. Including population growth leads to a critical threshold, such that for large flow/rheotaxis autoaggregation is suppressed. Non specified parameters are *δ* = 1 and *U* = 0.05).

For *ρ* > 0, however, NSCs have a bounded absolute maximum in *ν*^2^ ≥ 0. This implies a critical flow-rheotaxis threshold |*ωϕ*|* beyond which autoaggregation is not possible, see Fig. 7**B**. In this case, inclusion of rheotaxis can act to prevent clustering, even under strong chemotaxis scenarios.

The code used to implement the mathematical framework is available on request.
KJP was the sole author of this paper
The author declares that there are no conflicts of interest.
KJP is supported through the “MIUR—Dipartimento di Eccellenza” programme awarded to Dipartimento Interateneo di Scienze, Progetto e Politiche del Territorio.

## References

[1] Chapman JW, Klaassen RHG, Drake VA, Fossette S, Hays GC, Metcalfe JD, et al. Animal orientation strategies for movement in flows. Curr Biol. 2011;21(20):R861–R870.

[2] Stahl E. Zur Biologie der Myxomyceten. Botanische Zeitung. 1884;42:145–156.

[3] Lyon EP. On rheotropism. I.—Rheotropism in fishes. Am J Physiol. 1904;12(2):149–161.

[4] Bretherton FP, Rothschild NMV. Rheotaxis of spermatozoa. Proc Roy Soc London B Biol Sci. 1961;153(953):490–502.

[5] Hill J, Kalkanci O, McMurry JL, Koser H. Hydrodynamic surface interactions enable Escherichia coli to seek efficient routes to swim upstream. Phys Rev Lett. 2007;98(6):068101.

[6] Fu MHC, Powers TR, Stocker R. Bacterial rheotaxis. Proceedings of the National Academy of Sciences. 2012;109(13):4780–4785.

[7] Ricci N, Cionini K, Banchetti R, Erra F. Rheotaxis in Uronychia setigera (Ciliata, Hypotrichida). Journal of Eukaryotic Microbiology. 1999;46(3):268–277.

[8] Miki K, Clapham DE. Rheotaxis guides mammalian sperm. Current Biol. 2013;23(6):443–452.

[9] Mathewson RF, Hodgson ES. Klinotaxis and rheotaxis in orientation of sharks toward chemical stimuli. Comp Biochem Physiol A. 1972;42:79–82.

[10] Montgomery JC, Baker CF, Carton AG. The lateral line can mediate rheotaxis in fish. Nature. 1997;389(6654):960–963.

[11] Baker CF, Montgomery JC. The sensory basis of rheotaxis in the blind Mexican cave fish, Astyanax fasciatus. J Compar Physiol A. 1999;184(5):519–527.

[12] Gardiner JM, Atema J. Sharks need the lateral line to locate odor sources: rheotaxis and eddy chemotaxis. J Exp Biol. 2007;210(11):1925–1934.

[13] Suli A, Watson GM, Rubel EW, Raible DW. Rheotaxis in larval zebrafish is mediated by lateral line mechanosensory hair cells. PloS One. 2012;7(2).

[14] Olive R, Wolf S, Dubreuil A, Bormuth V, Debrégeas G, Candelier R. Rheotaxis of larval zebrafish: behavioral study of a multi-sensory process. Front Sys Neurosci. 2016;10:14.

[15] Oteiza P, Odstrcil I, Lauder G, Portugues R, Engert F. A novel mechanism for mechanosensory-based rheotaxis in larval zebrafish. Nature. 2017;547(7664):445–448.

[16] Rowat D, Gore M. Regional scale horizontal and local scale vertical movements of whale sharks in the Indian Ocean off Seychelles. Fisheries Research. 2007;84(1):32–40.

[17] Visser AW. Hydromechanical signals in the plankton. Marine Ecology Progress Series. 2001;222:1–24.

[18] Beauchamp RSA. Rheotaxis in Planaria alpina. Journal of Experimental Biology. 1933;10(2):113–129.

[19] Yuan J, Raizen DM, Bau HH. Propensity of undulatory swimmers, such as worms, to go against the flow. Proceedings of the National Academy of Sciences. 2015;112(12):3606–3611.

[20] Hamner WM, Hamner PP, Strand SW, Gilmer RW. Behavior of Antarctic krill, Euphausia superba: chemoreception, feeding, schooling, and molting. Science. 1983;220(4595):433–435.

[21] Fossette S, Gleiss AC, Chalumeau J, Bastian T, Armstrong CD, Vandenabeele S, et al. Current-oriented swimming by jellyfish and its role in bloom maintenance. Current Biology. 2015;25(3):342–347.

[22] Durand JP, Parzefall J. Comparative study of the rheotaxis in the cave salamander Proteus anguinus and his epigean relative Necturus maculosus (Proteidae, Urodela). Behavioural processes. 1987;15(2-3):285–291.

[23] Kobayashi DR, Farman R, Polovina JJ, Parker DM, Rice M, Balazs GH. “Going with the flow” or not: evidence of positive rheotaxis in oceanic juvenile loggerhead turtles (Caretta caretta) in the South Pacific Ocean Using Satellite Tags and Ocean Circulation Data. PLoS One. 2014;9(8).

[24] Parrish JK, Edelstein-Keshet L. Complexity, pattern, and evolutionary trade-offs in animal aggregation. Science. 1999;284(5411):99–101.

[25] Sumpter DJT. Collective animal behavior. Princeton University Press; 2010.

[26] Hay ME. Marine chemical ecology: chemical signals and cues structure marine populations, communities, and ecosystems. Ann Rev Mar Sci. 2009;1:193–212.

[27] Bonner JT. The social amoebae: the biology of cellular slime molds. Princeton University Press; 2009.

[28] Budrene EO, Berg HC. Complex patterns formed by motile cells of Escherichia coli. Nature. 1991;349:630–633.

[29] Reynierse JH, Gleason KK, Ottemann R. Mechanisms producing aggregations in planaria. Animal Behaviour. 1969;17:47–63.

[30] Walker KA, Jones TH, Fell RD. Pheromonal basis of aggregation in European earwig, Forficula auricularia L.(Dermaptera: Forficulidae). J Chem Ecol. 1993;19(9):2029–2038.

[31] Jumean Z, Gries R, Unruh T, Rowland E, Gries G. Identification of the larval aggregation pheromone of codling moth, Cydia pomonella. J Chem Ecol. 2005;31(4):911–924.

[32] Ratchford SG, Eggleston DB. Size-and scale-dependent chemical attraction contribute to an ontogenetic shift in sociality. Animal Behaviour. 1998;56(4):1027–1034.

[33] Lin CC, Prokop-Prigge KA, Preti G, Potter CJ. Food odors trigger Drosophila males to deposit a pheromone that guides aggregation and female oviposition decisions. Elife. 2015;4:e08688.

[34] Aggio J, Derby CD. Chemical communication in lobsters. In: Chemical communication in crustaceans. Springer; 2010. p. 239–256.

[35] Marquet N, Hubbard PC, da Silva JP, Afonso J, Canário AVM. Chemicals released by male sea cucumber mediate aggregation and spawning behaviours. Sci Rep. 2018;8(1):1–13.

[36] Hall MR, Kocot KM, Baughman KW, Fernandez-Valverde SL, Gauthier MEA, Hatleberg WL, et al. The crown-of-thorns starfish genome as a guide for biocontrol of this coral reef pest. Nature. 2017;544(7649):231–234.

[37] Kayal M, Vercelloni J, De Loma TL, Bosserelle P, Chancerelle Y, Geoffroy S, et al. Predator crown-of-thorns starfish (Acanthaster planci) outbreak, mass mortality of corals, and cascading effects on reef fish and benthic communities. PloS One. 2012;7(10):e47363.

[38] Keller EF, Segel LA. Initiation of slime mold aggregation viewed as an instability. J Theor Biol. 1970;26:399–415.

[39] Painter KJ. Mathematical models for chemotaxis and their applications in self-organisation phenomena. J Theor Biol. 2019;481:162–182.

[40] Hillen T, Painter K. Global existence for a parabolic chemotaxis model with prevention of overcrowding. Advances in Applied Mathematics. 2001;26(4):280–301.

[41] Domeier ML. In: Sadovy de Mitcheson Y, Colin PL, editors. Revisiting Spawning Aggregations: Definitions and Challenges. Springer Netherlands; 2012. p. 1–20.

[42] Jiang Y, Torrance L, Peichel CL, Bolnick DI. Differences in rheotactic responses contribute to divergent habitat use between parapatric lake and stream threespine stickleback. Evolution. 2015;69(9):2517–2524.

[43] Painter KJ, Hillen T. Spatio-temporal chaos in a chemotaxis model. Physica D. 2011;240:363–375.

[44] Wyatt TD. Pheromones and animal behaviour: Communication by smell and taste. Cambridge University Press; 2003.

[45] Flierl G, Grünbaum D, Levins S, Olson D. From individuals to aggregations: the interplay between behavior and physics. Journal of Theoretical biology. 1999;196(4):397–454.

[46] Chicoli A, Bak-Coleman J, Coombs S, Paley D. Rheotaxis performance increases with group size in a coupled phase model with sensory noise. Eur Phys J Spec Top. 2015;224(17):3233–3244.

[47] Pennington JT. The ecology of fertilization of echinoid eggs: the consequences of sperm dilution, adult aggregation, and synchronous spawning. Biological Bulletin. 1985;169(2):417–430.

[48] Crimaldi JP, Zimmer RK. The physics of broadcast spawning in benthic invertebrates. Ann Rev Mar Sci. 2014;6:141–165.

[49] Sadovy Y, Domeier M. Are aggregation-fisheries sustainable? Reef fish fisheries as a case study. Coral reefs. 2005;24(2):254–262.

[50] Hillesdon AJ, Pedley TJ, Kessler JO. The development of concentration gradients in a suspension of chemotactic bacteria. Bull Math Biol. 1995;57:299–344.

[51] Chertock A, Fellner K, Kurganov A, Lorz A, Markowich PA. Sinking, merging and stationary plumes in a coupled chemotaxis-fluid model: a high-resolution numerical approach. J Fluid Mech. 2012;694:155–190.

[52] Deleuze Y, Chiang CY, Thiriet M, Sheu TWH. Numerical study of plume patterns in a chemotaxis–diffusion–convection coupling system. Comp & Fluid. 2016;126:58–70.

[53] Chassignet EP, Hurlburt HE, Smedstad OM, Halliwell GR, Hogan PJ, Wallcraft AJ, et al. The HYCOM (hybrid coordinate ocean model) data assimilative system. Journal of Marine Systems. 2007;65(1-4):60–83.

[54] Murray JD. Mathematical Biology. II Spatial Models and Biomedical Applications. Springer-Verlag, New York; 2003.

[55] Satnoianu RA, Menzinger M. Non-Turing stationary patterns in flow-distributed oscillators with general diffusion and flow rates. Phys Rev E. 2000;62(1):113.

